# Human iPSC-derived microglia sense and dampen hyperexcitability of cortical neurons carrying the epilepsy-associated SCN2A-L1342P mutation

**DOI:** 10.1101/2023.10.26.563426

**Authors:** Zhefu Que, Maria I. Olivero-Acosta, Ian Chen, Jingliang Zhang, Kyle Wettschurack, Jiaxiang Wu, Tiange Xiao, C. Max Otterbacher, Muhan Wang, Hope Harlow, Ningren Cui, Xiaoling Chen, Brody Deming, Manasi Halurkar, Yuanrui Zhao, Jean-Christophe Rochet, Ranjie Xu, Amy L. Brewster, Long-jun Wu, Chongli Yuan, William C. Skarnes, Yang Yang

**Author notes:** these authors contributed equally to this work (Zhefu Que, Maria I. Olivero-Acosta). **To whom correspondence should be addressed:** Yang Yang, Ph.D.; Borch Department of Medicinal Chemistry and Molecular Pharmacology, College of Pharmacy, Purdue University, Hall for Discovery and Learning Research (DLR), 207 S Martin Jischke Dr, West Lafayette, IN 47907.

## Abstract

Neuronal hyperexcitability is a hallmark of seizures. It has been recently shown in rodent models of seizures that microglia, the brain’s resident immune cells, can respond to and modulate neuronal excitability. However, how human microglia interacts with human neurons to regulate hyperexcitability mediated by epilepsy-causing genetic mutation found in human patients remains unknown. The *SCN2A* genetic locus is responsible for encoding the voltage-gated sodium channel Nav1.2, recognized as one of the leading contributors to monogenic epilepsies. Previously, we demonstrated that the recurring Nav1.2-L1342P mutation identified in patients with epilepsy leads to hyperexcitability in a hiPSC-derived cortical neuron model from a male donor. While microglia play an important role in the brain, these cells originate from a different lineage (yolk sac) and thus are not naturally present in hiPSCs-derived neuronal culture. To study how microglia respond to diseased neurons and influence neuronal excitability, we established a co-culture model comprising hiPSC-derived neurons and microglia. We found that microglia display altered morphology with increased branch length and enhanced calcium signal when co-cultured with neurons carrying the Nav1.2-L1342P mutation. Moreover, the presence of microglia significantly lowers the action potential firing of neurons carrying the mutation. Interestingly, we further demonstrated that the current density of sodium channels in neurons carrying the epilepsy-associated mutation was reduced in the presence of microglia. Taken together, our work reveals a critical role of human iPSCs-derived microglia in sensing and dampening hyperexcitability mediated by an epilepsy-causing mutation present in human neurons, highlighting the importance of neuron-microglia interactions in human pathophysiology.

**Significance Statement:** Seizure studies in mouse models have highlighted the role of microglia in modulating neuronal activity, particularly in the promotion or suppression of seizures. However, a gap persists in comprehending the influence of human microglia on intrinsically hyperexcitable neurons carrying epilepsy-associated pathogenic mutations. This research addresses this gap by investigating human microglia and their impact on neuronal functions. Our findings demonstrate that microglia exhibit dynamic morphological alterations and calcium fluctuations in the presence of neurons carrying an epilepsy-associated SCN2A mutation. Furthermore, microglia suppressed the excitability of diseased hyperexcitable neurons, suggesting a potential beneficial role. This study underscores the role of microglia in the regulation of abnormal neuronal activity, providing insights into therapeutic strategies for neurological conditions associated with hyperexcitability.

## Introduction

Epilepsy is a neurological disorder characterized by recurrent and spontaneous seizures (Christensen et al., 2023); when left uncontrolled, it can lead to neuronal damage (Sun et al., 2022), cognitive deficits (Chai et al., 2023), and even sudden unexpected death (SUDEP) (Whitney et al., 2023). A prominent feature of epileptic seizures is hyperexcitability and excessive abnormal neural activity, which in some cases could be attributed to genetic changes in ion channels (Oyrer et al., 2018). The *SCN2A* gene encodes the pore-forming voltage-gated sodium channel alpha subunit 2, an ion channel protein that mediates neuronal action potential firing. *SCN2A* pathogenic heterozygous mutations are monogenic causes of epilepsies (Wolff et al., 2017; Yokoi et al., 2018; Epifanio et al., 2021; Yang et al., 2022; Zeng et al., 2022). In fact, a recent study positions *SCN2A* as the third most prevalent gene harboring epilepsy-related mutations (Knowles et al., 2022). Among many *SCN2A* disease-causing mutations, the *de novo* heterozygous missense mutation L1342P is a recurrent mutation associated with developmental and epileptic encephalopathy in multiple patients worldwide (Wolff et al., 2017; Crawford et al., 2021).

Previously, we developed an *in vitro* disease model of seizures with monolayer (2D) cortical neurons derived from human induced pluripotent stem cells (hiPSCs) carrying the epilepsy-associated Nav1.2-L1342P mutation. Our investigation revealed intrinsic and network neuronal hyperexcitability in the hiPSC-derived cortical neurons carrying this Nav1.2-L1342P mutation (Que et al., 2021). While neuronal hyperexcitability appears to be an intrinsic property of diseased neurons, emerging evidence suggests that non-neuronal cell types can influence neuronal excitability and thus can contribute to disease severity or progression (Chen et al., 2023). Microglia are brain-resident macrophages that originate from the yolk sac and migrate into the brain during development (Speicher et al., 2019). Mounting evidence suggests that microglia play a key role in regulating brain homeostasis and maintaining neural circuit integrity (Olson and Miller, 2004; Paolicelli et al., 2011; Schafer et al., 2012; McQuade et al., 2018; Weinhard et al., 2018). The impact of microglia on neuronal excitability has been extensively studied in rodents and found to be multifaceted. For instance, in certain cases, activated microglia release pro-inflammatory mediators that enhance neuronal activity, thereby promoting epileptogenesis (Henning et al., 2023). However, in other studies, microglia exert a regulatory role in limiting excessive neuronal activity, as observed in rodent models of chemically induced seizures (Eyo et al., 2014; Badimon et al., 2020; Merlini et al., 2021). While these published studies are vital for us to understand microglia-neuron interactions, these studies rely on rodent models in which seizures and neuronal hyperexcitability are triggered by external chemicals (Eyo et al., 2014; Liu et al., 2020). No studies, however, have yet reported how human microglia respond to and regulate the excitability of intrinsically hyperexcitable human neurons carrying epilepsy-causing genetic mutations, hindering our understanding of microglia in human disease conditions.

In the current study, we used a co-culture system of hiPSC-derived microglia and hiPSCs-derived cortical neurons carrying the epilepsy-associated Nav1.2-L1342P mutation to study microglia-neuron interactions. We found an increase in the length of microglial branches (processes) when co-cultured in the presence of the mutant Nav1.2-L1342P neurons. This effect paralleled an increase in calcium signaling within microglial processes. In addition, we demonstrated that hiPSC-derived microglia reduced hyperexcitability and sodium current density in neurons carrying the Nav1.2-L1342P genetic mutation. Taken together, our study illustrates the vital role of microglia in human epilepsy pathophysiology. Our study also suggests a complex microglial-neuronal interaction with both cell types influencing each other’s phenotypes.

## Materials and Methods

### Generation of hiPSC-derived cortical neurons

Previously described human induced pluripotent stem cells (hiPSCs) from a male donor expressing wild-type (Control) and CRISPR/Cas9-engineered Nav1.2-L1342P mutant channels (Que et al., 2021) were used in this study. Experiments were performed with four human iPSC lines (WT: KOLF2.1 and A03; L1342P: A12 and E09). Feeder-free hiPSCs colonies were grown on Matrigel (Corning, Catalog No. 354230) and maintained in StemFlex (ThermoFisher, Catalog No. A3349401) with daily media changes. Quality controls were performed for all lines, including Sanger sequencing, karyotyping, and immunocytochemistry. Undifferentiated hiPSC colonies displayed normal and homogenous morphology with defined edges and low levels of spontaneous differentiation. In addition, they consistently expressed standard pluripotency markers, including SOX2, TRA-1-80, OCT4, TRA-1-60, NANOG, and SSEA1 (data not shown).

A dual-SMAD inhibition method utilizing embryoid bodies (EBs) was used to generate cortical neurons based on our established protocol (Que et al., 2021). To generate embryoid bodies (EBs), hiPSC colonies were dissociated into single cells with Accutase (Innovative Cell Technologies, Catalog No.AT104) and seeded with 10 mM rock inhibitor (RevitaCell Supplement, Invitrogen, Catalog No. A2644501) for the initial 24 hours on ultra-low attachment 96-well plates (Corning, Catalog No. CLS3474-24EA) with a cell density of ∼12,000 cells per microwell in an EB formation medium containing neural induction medium (StemCell Technologies, Catalog No. 05835) supplemented with 100 nM of LDN-193189 (Sigma, Catalog No. SML0559) and 10 μM SB431542 (Tocris, Catalog No. 1614) to begin DUAL SMADi neural induction. After seven days, EBs were collected and seeded on Matrigel until the appearance of neural rosettes. A neural rosette selection reagent (StemCell Technologies, Catalog No. 05832) was used to lift the rosette monolayer clusters, which were dissociated and seeded onto Matrigel-coated plates until the appearance of neural progenitor cells (NPCs). Stocks of NPC were frozen and stored for later differentiations.

To begin neural differentiation, neural progenitors were plated on poly-L-ornithine (PLO)-laminin-coated vessels at a density of ∼2.5x10^4^ cells/cm^2^ and differentiated for about three weeks in a media containing Neurobasal Plus medium (Invitrogen, Catalog No. A3582901), 1X Non-Essential Amino Acids solution (NEAA; Invitrogen, Catalog No. 11140050), 1X GlutaMAX (Invitrogen, Catalog No. 3505006), PenStrep (10,000 U/mL; Gibco, Catalog No. 15-140-163), 1X B27 plus supplement (Gibco, Catalog No. A3582801), 1X N2 Supplement (Gibco, Catalog No.17-502-048), 100 μm dibutyryl cAMP (dcAMP; Santa Cruz Biotechnology, Catalog No. sc-201567A), 200 μm ascorbic acid (Wako Chemicals; Catalog No. 323-44822), 20 ng/mL brain-derived neurotrophic factor (BDNF; ProspecBio, Catalog No. CYT-207), 20 ng/mL glial cell-derived neurotrophic factor (GDNF; ProspecBio, Catalog No. CYT-305) with media replacement every 2–3 days. After 20 days, cells were replated at 1.5x10^4^ cells/cm^2^ on glass coverslips using a complete maturation media formulation containing Neurobasal Plus (Gibco, Catalog No. A3582901). After a week, the basal media was changed into Brainphys (StemCell Technologies, Catalog No. 05790), PenStrep, 1X N2 supplement, 1X B27 plus supplemented with BDNF, GDNF, and cAMP in the same concentration previously described.

### Generation of hiPSC-derived microglia

To produce hiPSC-derived microglia, we used two different control (WT) human iPSC lines, KOLF2.1 and GCaMP6f-H04. The KOLF2.1 cell line-derived microglia were co-cultured with neurons for electrophysiology experiments. The GCaMP6f-H04 line has the GCaMP6f calcium indicator (Chen et al., 2013) engineered into the AAVS1 safe harbor locus of the reference Kolf2.1 iPSCs line using CRISPR/Cas9 mediated knock-in and was used for the microglial calcium imaging. Specifically, a commercially available plasmid (pAAVS1-PC-GCaMP6f, plasmid #73503) containing the GCaMP6f knock-in construct was used to generate the GCaMP6f-H04 line from KOLF2.1 reference iPSCs line. For each experiment, at least two differentiations (biological replicates) were performed for each cell line.

To begin microglia differentiation, we largely followed a commercially available kit based on well-established, previously published protocols (Abud et al., 2017; McQuade et al., 2018; McQuade and Blurton-Jones, 2021). Briefly, feeder-free hiPSCs were guided towards a mesodermal, hematopoietic lineage to obtain hematopoietic progenitor cells (HPCs) using the STEMdiff Hematopoietic Kit (StemCell Technologies, Catalog No. 05310). Next, HPCs were converted into homeostatic microglia with a differentiation medium composed of DMEM/F12 basal media (Gibco, Catalog No. 11-320-033), 2X 100X Insulin-Transferrin-Selenium (ITS-G)(Gibco, Catalog No. 41400045), 2X B27, 0.5X N2, 1X GlutaMAX (Gibco, Catalog No. 35050061), 1X non-essential amino acids (Gibco, Catalog No. 11140050), 400 µM monothioglycerol (Sigma, Catalog No. M6145-25ML), 5 µg/mL insulin (Sigma, Catalog No. I2643-25MG). Before use, this media was supplemented with 25 ng/mL cytokine M-CSF (Macrophage colony-stimulating factor 1) (Sigma, Catalog No. 300-25), 100 ng/mL IL-34 (Interleukin-34) (Peprotech, Catalog No. 200-34), and 50 ng/mL TGFβ-1 (Transforming growth factor beta 1) (Peprotech, Catalog No. 100-21) until day 24. Then, cells were cultured in a maturation medium with the same composition as the differentiation medium, with the addition of 100 ng/mL of CD200 (Cluster of Differentiation 200/OX-2 membrane glycoprotein) (ACROBiosystems, Catalog No. 50-101-8369) and 100 ng/mL of CX3CL1 (Fractalkine/ chemokine (C-X3-C motif) ligand 1) (Peprotech, Catalog No. 300-31) for up to 12 days.

### Co-culture of hiPSC-derived microglia and hiPSC-derived cortical neurons

As microglia developmentally come from the yolk sac and thus do not naturally exist in hiPSCs-derived neuronal culture, a co-culture model must be established. To achieve this, hiPSC-derived microglia were generated and seeded on cortical neuron monolayers, which had been differentiated for a minimum of 38 days. The co-cultures were set up in a 96-well clear-bottom plate at a density of 10,000 cells per well, maintaining a 1:1 ratio between microglia and neurons. The co-cultures were incubated for up to seven days to allow for the formation of interactions between the two cell types. For medium exchange, a mixture of half-complete neurobasal medium and half microglia maturation medium was used, with media exchanges performed every two days.

### Immunocytochemistry

hiPSC-derived microglia cells were cultured on glass coverslips (Neuvitro, Catalog No. GG-12-Pre) or 24-well glass-bottom plates with #1.5 cover glass (Celvis, Catalog No. P24-1.5H-N) previously coated with a 1:5 mixture of poly-L-ornithine (PLO) and phosphate-buffered saline (PBS)-Laminin. Before the experiment, the samples were briefly washed in PBS (Corning, Catalog No. 21-040-CMX12) and then fixed in 4% paraformaldehyde in PBS at room temperature (RT) for 15 minutes. Following fixation, the samples were rinsed three times with PBS (5 minutes per rinse) and permeabilized with 0.3% Triton X-100 (pH 7.4) surfactant for 20 minutes.

The samples were treated with 5% bovine serum albumin (BSA; Sigma Catalog No. 9048) to block nonspecific binding for one hour at RT. Subsequently, the samples were incubated overnight at 4°C in a humidified chamber with primary antibodies diluted in 1% BSA. The primary antibodies used included rabbit anti-P2RY12 (Purinergic Receptor P2Y12; Sigma Prestige Antibodies, Catalog No. HPA014515), rabbit anti-TMEM119 (Transmembrane Protein 119; Sigma Prestige Antibodies, Catalog No. HPA051870), and rabbit anti-IBA1 (Ionized calcium-binding adaptor molecule 1; Abcam, Catalog No. 178846). The following day, the samples were rinsed three times with PBS and then incubated with fluorescent-dye-conjugated secondary antibodies, which were diluted in 1% BSA for 2 hours at room temperature (RT) in the dark. After the incubation, the secondary antibody solution was removed, and the coverslips were washed three times with PBS (5 minutes per wash) in the dark. The secondary antibodies used for hiPSC-derived neurons and microglia were as follows: anti-rabbit or anti-mouse antibodies conjugated with Alexa Fluor 488 (Invitrogen, 1:1000), anti-rabbit or anti-mouse antibodies conjugated with Alexa Fluor 555 (Invitrogen, 1:1000), anti-guinea pig antibodies conjugated with Alexa Fluor 488 (Invitrogen, Catalog No. A11073, 1:1000), and anti-rat antibodies conjugated with Alexa Fluor 647 (Invitrogen, Catalog No. A21247, 1:1000). For DAPI counterstaining, either VECTASHIELD antifade mounting medium with DAPI (Vector Laboratories, Catalog No. H-1200) or a PBS-DAPI solution (ThermoFisher, Catalog No. 62238, 1:10,000) was used. For staining hiPSC-derived neurons, primary antibodies for mouse anti-MAP2 (microtubule-associated protein 2; Invitrogen, Catalog No. 13-1500, 1:1000) and guinea pig anti-synapsin1/2 (Synaptic Systems, Catalog No. 106044, 1:1000) were used. Microglial images were acquired with a Nikon Ti2 Eclipse fluorescence microscope, while images of the microglia-neuron co-culture were captured using a ZEISS LSM 900 Airy Scan microscope.

### Incucyte SX5 Live-Cell phagocytotic assay

To assess the phagocytic activity of microglia derived from hiPSCs, we used pHrodo-Myelin, a pH-sensitive dye combined with a myelin fragment (Hendrickx, Schuurman, van Draanen, Hamann, & Huitinga, 2014), graciously provided by Dr. Shaoyou Chu from Indiana University (Mason, Soni, & Chu, 2023). First, we prepared a Corning #1.5 glass-bottom 96-well by coating it with a 5X diluted solution of Poly-D-Lysine (PDL) in 1x PBS, letting it incubate at 37°C overnight. Subsequently, the wells were washed three times with 1x PBS. Approximately 15,000 matured hiPSC-derived microglia cells, suspended in 100 μL of microglia maturation media, were placed delicately onto the coated wells. The plate was then left in an incubator at 37°C overnight. The following day, we prepared a stock of pHrodo-myelin at 5 μg/mL in the microglia maturation media. 50 μL of the media was extracted from each well, and 50 μL of the pHrodo-myelin stock was added, resulting in a final concentration of 2.5 μg/mL of phrodo-myelin. To monitor the uptake of the pHrodo-myelin dye for 48 hours, we employed an Incucyte SX5 imaging system with an Orange Filter and Brightfield using a 20X objective. The exposure time was maintained at 300 ms. To quantify the phagocytic activity, we calculated the normalized integrated intensity of pHrodo-myelin by dividing the total integrated intensity of pHrodo-myelin corresponding to each well over the area occupied by the microglial cells, as identified by the brightfield channel.

### Morphological Characterization of hiPSC-derived IBA1+ Microglia

To assess alterations in microglia morphology following co-culture with control (WT) and Nav1.2-L1342P cortical neurons, we conducted immunocytochemistry staining for microglia-specific marker IBA1. The ImageJ Analyze Skeleton plugin (available at https://imagej.net/plugins/analyze-skeleton/) was employed to generate a skeletonized representation of microglial morphology, which was used to determine the unique average length of the microglial processes per field of view (Jairaman et al., 2022). To obtain area, circularity, and perimeter measurements, we manually selected clearly defined microglial morphologies using the NIS-elements ROI editor and used the automatic measurement results to obtain values. We measured between 7-35 microglia per field of view.

### Live-cell calcium imaging of hiPSC-derived microglia expressing GCaMP6f

Live cell calcium imaging was conducted using an inverted widefield Nikon Eclipse Ti2 microscope. Time-lapse recordings were acquired at a frequency of 1 Hz for 200 seconds. Microglial somata and processes were manually defined using the FIJI’s ImageJ (Version 2.3.0/1.53f) (Schindelin et al., 2012) software’s ROI (Region of Interest) editor. Time measurements were performed to detect the spontaneous calcium transient fluctuations. The signal intensity was normalized as ΔF/F, where ΔF represents the difference between an individual ROI’s fluorescence intensity (F) and its minimum intensity value (Fmin). The normalized data was then transferred to OriginPro (Origin2021b version 9.8.5.201) for peak detection and quantifications. We used the Nikon Imaging Software Elements (NIS Elements Version 5.02) for representative image processing.

### Electrophysiology of hiPSC-derived neurons co-cultured with hiPSC-derived microglia

The experiment involved performing whole-cell patch-clamp recordings using an EPC10 amplifier and Patchmaster v2X90.3 software (HEKA Elektronik) paired to an inverted microscope configuration (NikonTi-2 Eclipse). For experiments under the voltage clamp configuration, we used our previously reported patch solution and protocol (Que et al., 2021). Thick-wall borosilicate glass pipettes (BF150-86-10) were pulled to reach the resistances of 2–4 MΩ. Briefly, the activation curve from the voltage-gated sodium channel was achieved by 10 ms steps from -70 to +50 mV in a 5 mV increment, with a holding potential of −100 mV. The currents from both groups were recorded at 5 min after obtaining the whole-cell configuration. P/N leak subtraction procedure was applied during the protocol. The current density value was obtained using the current under each voltage command divided by the capacitance of the neuron. For the current-clamp recording, the external solution contained the following: 140 mM NaCl, 5 mM KCl, 2 mM CaCl_2_, 2 mM MgCl_2_, 10 mM HEPES, and 10 mm dextrose, titrated with NaOH to pH 7.3. The internal solution contained the following: 128 mM K-gluconate, 5 mM KCl, 5 mM NaCl, 1 mM MgCl2, 3 mM Mg-ATP, 1 mM EGTA, 10 mM HEPES, and 10 mM dextrose, titrated with KOH to pH 7.2. The osmolarity was brought to 320 and 310 mOsm by adding dextrose for the extracellular and internal solutions, respectively. The glass pipettes (BF150-86-10) were used and pulled to reach the resistances of 4-8 MΩ. We measured the repetitive action potential (AP) firings at increased current injections using a prolonged 800-ms present stimulus ranging from 0 to 125 pA in 5-pA increments. Neurons were transduced with the AAV-CamKII-GFP virus (Addgene 50469-AAV9), and neurons with GFP-positive signals were selected for patch clamp experiments.

### Statistical analysis

GraphPad Prism (version 9.5.1) and OriginPro 2021b (version 9.8.5.201) were used for statistical analysis. The number of experimental samples (n) in each group was indicated in the figure legend. Results are presented as mean ± standard error of the mean (s.e.m). Values are shown in figures as *p < 0.05, **p < 0.01, ***p < 0.001, ****p < 0.0001, and n.s. (not significant) as p ≥ 0.05. The detailed statistical methods were outlined in the figure legends.

## Results

### hiPSC-derived microglia express relevant markers and display phagocytotic capacity

While microglia play an indispensable role in the brain, they developmentally originate from the yolk sac and thus do not naturally appear in hiPSC-derived neuronal culture (Lukens and Eyo, 2022); therefore, a co-culture model of neurons and microglia needs to be established to study the interaction between these cells. To this end, neurons and microglia were differentiated using distinct protocols and cultured together. To generate glutamatergic cortical neurons, we differentiated hiPSCs using our published protocol (Que et al., 2021) **(Figure 1A)**, which can yield electrically active matured neurons in 45 days *in vitro* (post-neural progenitor cell stage). These neurons express synaptic and neuron-specific markers such as Synapsin1/2 (SYN1/2) and Microtubule-associated protein-2 (MAP2) **(Figure 1B)**. Since Nav1.2 is minimally expressed in microglia from mice and human iPSC-derived microglia (Black et al., 2009; Black and Waxman, 2012, 2013; Abud et al., 2017; Grubman et al., 2020; Dräger et al., 2022), we intentionally used microglia derived from control reference (Wild-type/WT) hiPSC lines to simplify the research design. Control (Wild-type/WT) mature microglia were generated after 24 days of differentiation **(Figure 1C)** and were characterized via staining and quantification of microglia-specific markers, including IBA1, TMEM119, and P2RY12 **(Figure 1D)**. We revealed a high percentage of microglial cells after differentiation (98.43 ± 0.44% TMEM119 labeling, n=18 fields of view, two differentiations; 97.18 ± 1.44% IBA1 labeling, n=13 fields of view; two differentiations; and 93.59± 0.45 % of P2RY12 labeling, n=19 fields of view, two differentiations). Our results thus indicate that hiPSC-derived microglia can be successfully obtained in a relatively homogeneous population. Next, we investigated the phagocytic function of the differentiated microglia. As immune and professional phagocyte cells, microglia can engulf substrates like synaptic and myelin debris. To evaluate the phagocytic capacity of the mature microglia, we exposed them to pHrodo-labeled myelin particles. Over 36 hours, we observed the continuous internalization of myelin by the microglia, an ability that was also demonstrated in primary human microglia (Hendrickx et al., 2014), indicating their robust phagocytic ability **(Figure 1E)**.

**Figure 1.**
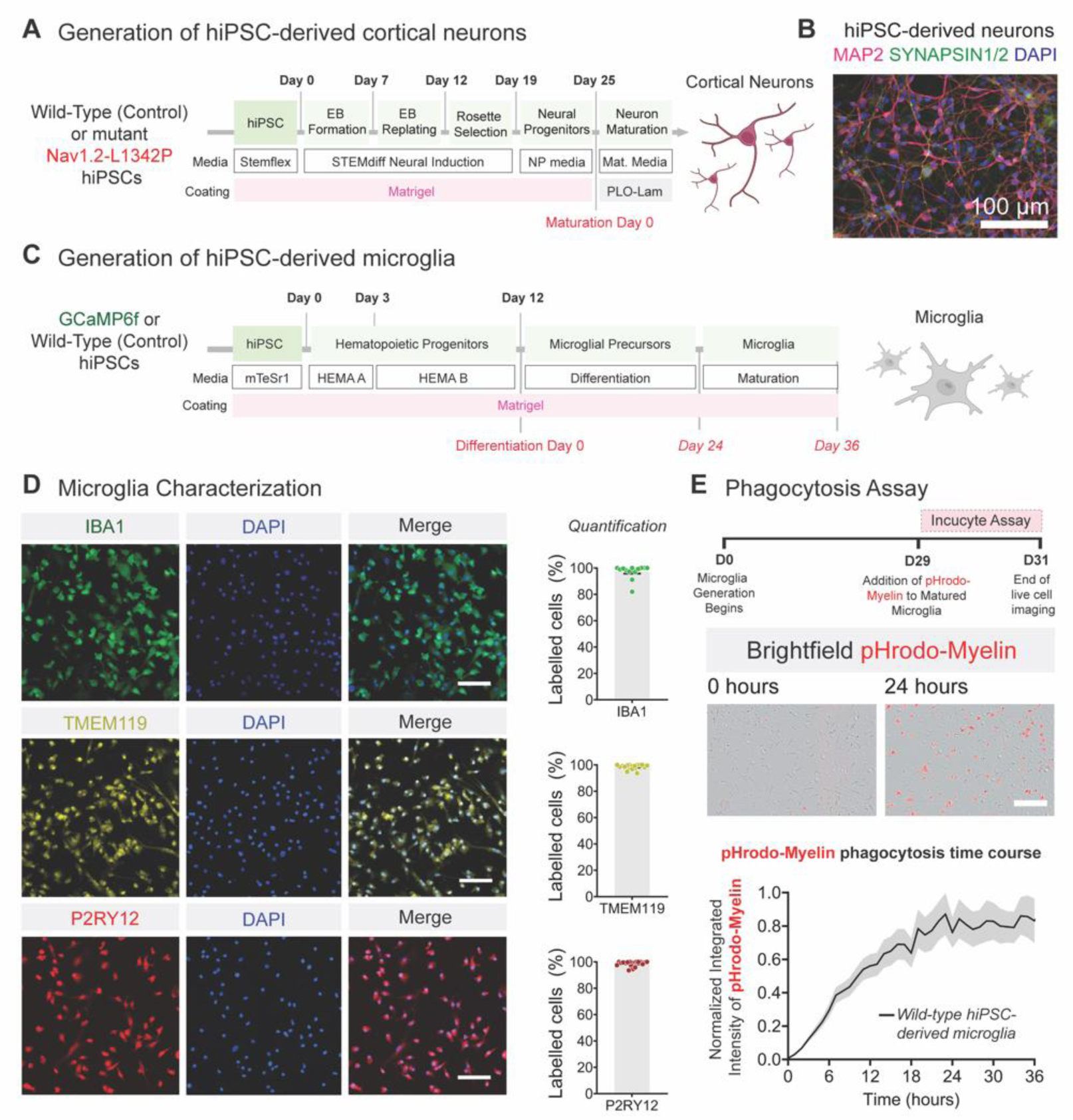
Characterization of hiPSC-derived cortical neurons and microglia. **(A)** Schematic illustrating the protocol for generating hiPSC-derived cortical neurons. **(B)** Representative Fluorescent image of hiPSC-derived neurons stained for somatodendritic marker MAP2 (magenta), synaptic vesicle proteins SYN1/2 (green), and DAPI (blue). **(C)** Schematic illustrating the protocol for generating hiPSC-derived microglia. Human iPSCs are differentiated into hematopoietic progenitor cells for 12 days and cultured in microglia differentiation media for 24 days. The microglia maturation process is then carried out for up to 12 days. **(D)** Representative images of hiPSC-differentiated microglia expressing microglial-specific markers: IBA1 (D, top panel, green, n=13 fields of view, two differentiations), TMEM119 (D, middle panel, yellow, n=18 fields of view, two differentiations), P2RY12 (D, lower panel red, n=19 fields of view, two differentiations). DAPI was used to stain nuclei. Data is presented as mean ± s.e.m. Scale bar=100 μm. **(E)** Phagocytosis of pHrodo-myelin by wild-type (control) hiPSC-derived microglia. Data was obtained from one differentiation of three wells (48 images per well). Representative images at 0 hours and 24 hours after the addition of pHrodo-myelin. Human iPSC-derived microglia phagocytosed the pHrodo-labeled bioparticles, showing a gradually increasing red fluorescent signal over time. Scale bar=25 μm. hiPSC, human induced pluripotent stem cells; EB, Embryoid body; NP, Neural Progenitors; MAP2, microtubule-associated protein-2; SYN1/2, Synapsin1/2; IBA1, Ionized calcium-binding adaptor molecule 1; TMEM119, transmembrane protein 119; P2RY12, Purinergic Receptor P2Y12.

### Microglia co-cultured with Nav1.2-L1342P cortical neurons display morphological alterations

In rodent models, microglia extend and retract their extended processes to survey the brain and regulate network hyperexcitability (Merlini et al., 2021). However, how human microglia respond to human neurons carrying seizure-related genetic mutations is not known. Thus, we sought to investigate whether hiPSC-derived microglia would exhibit morphological changes when co-cultured with hiPSC-derived hyperexcitable neurons carrying a Nav1.2-L1342P mutation (Que et al., 2021). To achieve this, we conducted a co-culture experiment by seeding the microglia on top of neuronal monolayers and maintaining them for at least seven days, as illustrated in **Figure 2A**. We confirmed the presence of microglia and neurons in the co-cultures with IBA1 (microglia; red) and MAP2 (neurons, green) via immunocytochemistry (**Figure 2B)**.

**Figure 2.**
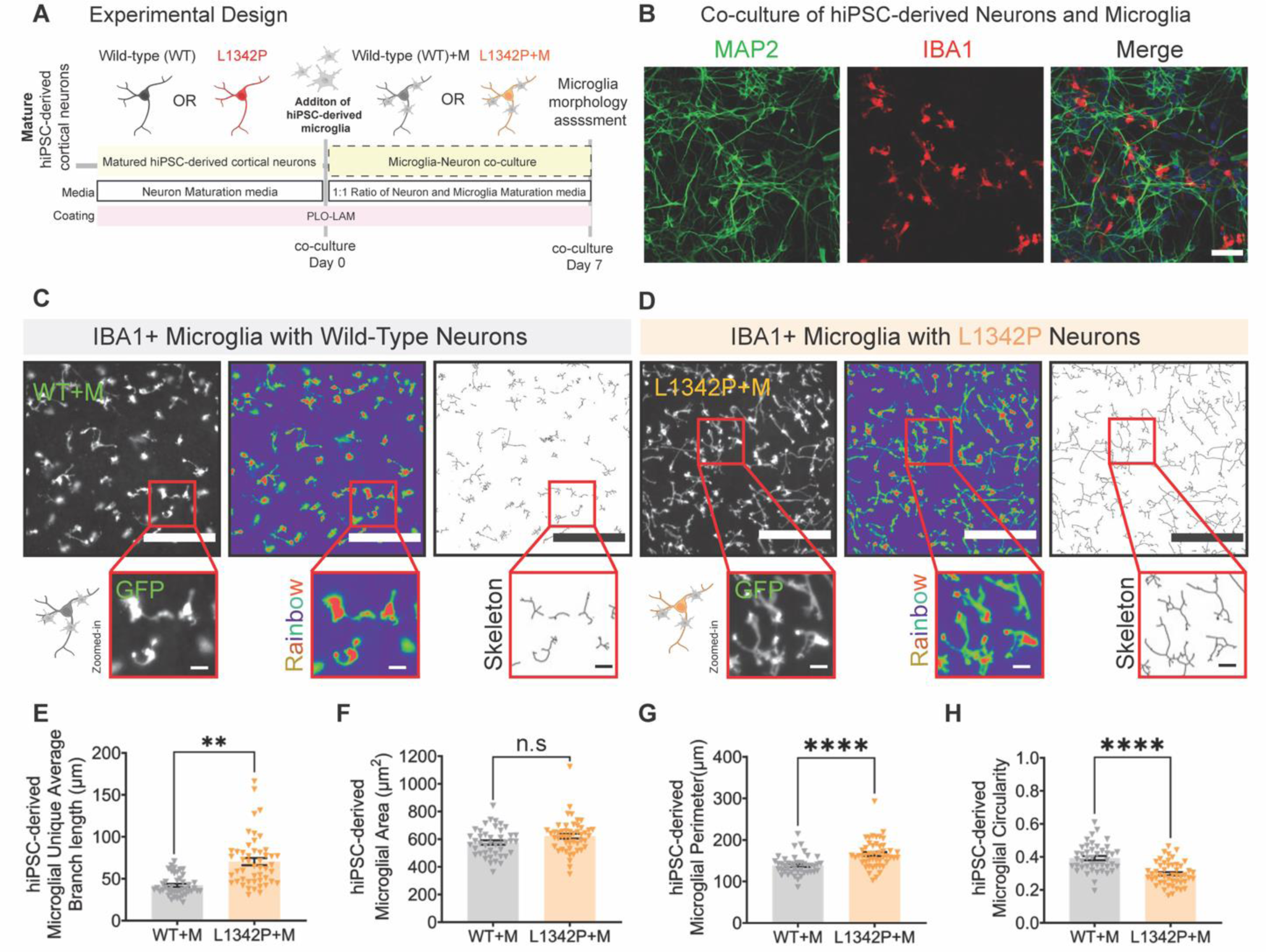
Human microglia in co-culture with L1342P neurons display morphological changes. **(A)** The hiPSC-derived neurons and microglia were matured separately, and then microglia were seeded on top of neurons for seven days before imaging. **(B)** Representative images of co-cultured neurons stained for neuron-specific marker MAP2 (green), microglia stained for IBA1 (red), and DAPI (blue) as a nuclear stain**. (C-D)** Human iPSC-derived microglia in co-culture with control (WT) neurons (WT+M, **C**) and Nav1.2-L1342P neurons (L1342P+M, **D**). IBA1+ microglia co-cultured with the hyperexcitable Nav1.2-L1342P neurons displayed an extended ramified process compared to co-culture with control (WT) neurons. Images are pseudo-colored in a rainbow gradient to facilitate identification, and the skeletonized view was included to detail branches. **(E)** The microglial average branch length increases in co-culture with Nav1.2-L1342P cortical neurons (WT+M: n=43 fields of view and L1342+M: n = 49 fields of view, three differentiations, two clones per condition). **(F)** The total microglial area did not change in co-culture with Nav1.2-L1342P neurons (WT+M: n=43 fields of view and L1342+M: n=49 fields of view, three differentiations, two clones per condition). **(G)** Microglial perimeter is enhanced in co-culture with Nav1.2-L1234P neurons, indicating extended processes (WT+M: n=43 fields of view and L1342+M: n=49 fields of view, three differentiations, two clones per condition). **(H)** Microglial circularity is decreased in co-culture with Nav1.2-L1342P neurons, indicating that they are less ameboid-like (WT+M: n=43 fields of view, and L1342+M: n=49 fields of view, three differentiations, two clones per condition). Each dot represents the mean value of a parameter per field of view. Data are presented as mean ± s.e.m. Scale bar=50 μm. Data was pooled from three differentiations. Data was analyzed by nested *t-test*; **p < 0.01 and ****p < 0.0001.

We traced the IBA1-positive microglial processes in co-culture with hiPSC-derived cortical neurons **(Figure 2C-D)**. Our results revealed that microglia exhibited a significant increase in process length per cell when co-cultured with L1342P mutant neurons versus when co-cultured with control (WT) neurons (WT+M: 42.08 ± 1.89 μm, n=43 fields of view, L1342P+M: 70.40 ± 4.27 μm, n=49 fields of view, three differentiations, two clones per condition, \*\**p*<0.01, Nested *t-test*, **Figure 2E**). In terms of total microglial area (soma and processes), we found no significant alterations in either co-culture states, indicating there were no changes in microglial cell size (WT+M: 575.30 ± 15.30 μm^2^, n=43 fields of view, L1342P+M: 621.50 ± 17.21 μm^2^, n=49 fields of view, three differentiations, two clones per condition, non-significant, Nested *t-test*, **Figure 2F**). However, the microglia perimeter was increased in microglia co-cultured with the Nav1.2-L1342P cortical neurons. Considering microglia size did not change, this data could be interpreted as microglial processes, such as extensions and branches being altered (WT+M: 138.50 ± 3.56 μm, n=43 fields of view, L1342P+M: 166.00 ± 4.67 μm, n=49 fields of view, three differentiations, two clones per condition, \*\*\*\**p*<0.0001, Nested *t-test*, **Figure 2G**). Microglia circularity was also calculated (circularity index is a number between 0-1, and 1 represents a perfect round shape). We found that the microglia circularity index was also decreased when co-cultured with the Nav1.2-L1342P neurons (WT+M: 0.3924 ± 0.01 μm, n=43 fields of view, L1342P+M: 0.30 ± 0.01 μm, n=49 fields of view, three differentiations, two clones per condition; \*\*\*\**p*<0.0001, *Nested t-test*, **Figure 2H**). Taken together, our data showed that microglia co-cultured with mutant Nav1.2-L1342P neurons display altered morphological changes compared to microglia co-cultured with control (WT) neurons, revealing that hyperexcitable neurons influence the morphology of microglia.

### Calcium signaling in hiPSC-derived microglia is enhanced when co-culture with L1342P neurons

Previous work using *in vivo* mouse models indicated that microglial calcium signals could respond to chemically triggered neuronal activity changes (Umpierre et al., 2020). However, how human microglia sense and respond to intrinsically hyperexcitable human neurons carrying an epilepsy mutation remains to be elucidated. Thus, we carried out experiments to address this gap. To study the calcium dynamics of our hiPSC-derived microglia, we established a hiPSC line in which the endogenous AAVS1 safe harbor site was engineered with a GCaMP6f gene using CRISPR/Cas9. We differentiated the GCaMP6f-iPSC lines into microglia and co-culture them with hiPSC-derived cortical neurons. Our results demonstrated that GCaMP6f hiPSC-derived microglia in co-culture with either WT neurons or neurons carrying Nav1.2-L1342P mutation can display spontaneous Ca^2+^ activity, reflected in continuously changing GCaMP6f calcium signal fluctuations (**Figure 3A-B).** This assay thus allows us to determine how calcium dynamics in microglia respond to neurons with different excitability.

**Figure 3.**
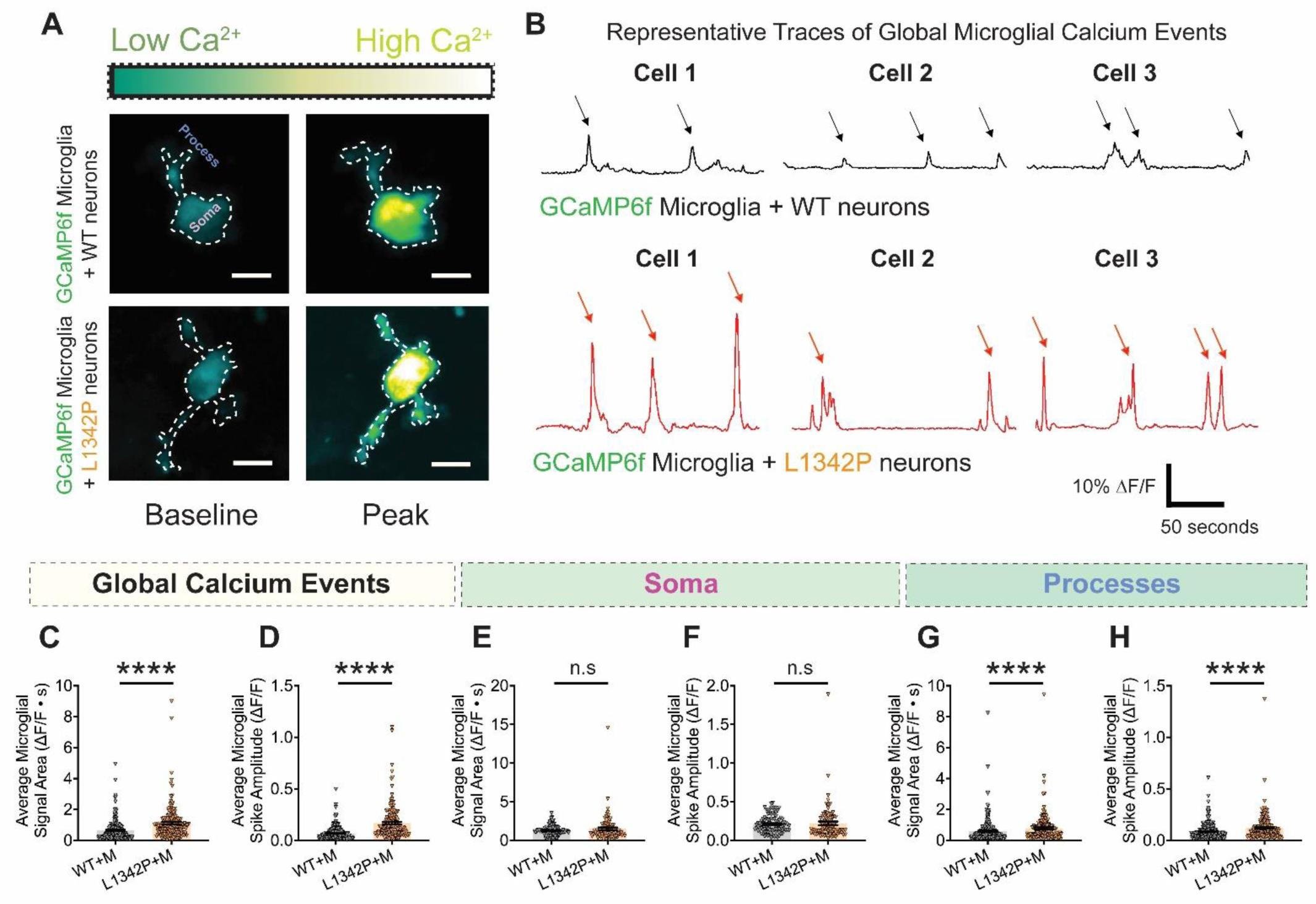
GCaMP6f calcium signal is enhanced in human microglia processes when co-culturedwith hyperexcitable hiPSC-derived L1342P neurons. **(A)** Fluorescent images of hiPSC-derived microglia expressing GCaMP6f co-cultured with WT (top) and mutant Nav1.2-L1342P (bottom) hiPSC-derived neurons. The GCaMP6f calcium signal is pseudocolored, with dark green indicating low signal and yellow hues depicting high signal. **(B)** Representative ΔF/F traces of GCaMP6f global calcium activity of microglia in co-culture with control (WT) neurons (top, black) or Nav1.2-L1342P neurons (bottom, red). Three representative cells per condition are shown. **(C)** The global average microglia calcium signal area increases in co-culture with Nav1.2-L1342P neurons. **(D)** The global average microglial calcium spike amplitude increases in co-culture with Nav1.2-L1342P neurons. **(E)** The average microglial signal area in the soma microdomain is not statistically different in the two co-culture conditions. **(F)** The average microglial spike amplitude in the soma microdomain is not statistically different in the two co-culture conditions. **(G)** The average microglial calcium signal area of the processes microdomain is increased in co-culture with Nav1.2-L1342P neurons. **(H)** The processes’ average microglial calcium spike amplitude increases in co-culture with L1342P neurons. Data was collected from two independent differentiations per genotype. Data in C, D, E, F, G, and H were analyzed with *Mann-Whitney’s U test*. ****p < 0.0001, and n.s (not significant).

Microglia are made up of microdomains (soma and extended processes) that could have different calcium signaling patterns (Umpierre et al., 2020). Therefore, we divided calcium fluctuation measurements into three categories: **1)** entire (global) microglial calcium activity and compartmentalized activity within **2)** soma, and **3)** processes. We first analyzed the calcium activity patterns from the entire microglia compartment. We found that the spikes generated from microglia in co-culture with Nav1.2-L1342P neurons had a larger amplitude than those found in microglia co-cultured with control (WT) neurons when the measurement was done for the entire microglial area. Specifically, we observed an elevated calcium signal area under the curve (AUC) in microglia co-cultured with Nav1.2-L1342P neurons, almost twice higher than than that observed in microglia co-cultured with WT neurons (WT+M: 0.652 ± 0.045, n=209 cells and L1342P+M: 1.125 ± 0.086, n=168 cells, ****p<0.0001, *Mann-Whitney U test*, **Figure 3C**). Similar to the increased AUC, we observed that the average microglial spike amplitude increased more than two fold when the microglia were co-cultured with Nav1.2-L1342P neurons (WT+M: 0.069 ± 0.004, n=204 cells and L1342P+M: 0.168 ± 0.012, n=168 cells, ****p<0.0001, *Mann-Whitney U test*, **Figure 3D**).

When analyzing the calcium dynamics of separate microglial sub-compartments, we found no significant difference in the average calcium signal AUC for the soma (WT+M: 1.238 ± 0.066, n=100 cells, L1342P+M: 1.494 ± 0.175, n=92 cells, ns *p = 0.987*, *Mann-Whitney U test*, **Figure 3E**). There was also no significant difference in the average microglial spike amplitude per soma between microglia co-cultured with either WT neurons or Nav1.2-L1342P neurons (WT+M: 0.211 ± 0.011, n=100 cells, L1342P+M: 0.221 ± 0.023, n=92 cells, ns *p= 0.281*, *Mann-Whitney U test*, **Figure 3F**). On the other hand, we found that the average calcium signal AUC for the processes was significantly higher in microglia co-cultured with Nav1.2-L1342P neurons compared to these in microglia co-cultured with control (WT) neurons (WT+M: 0.595 ± 0.047, n=250 cells, L1342P+M: 0.786 ± 0.067, n=183 cells, ****p<0.0001, *Mann-Whitney U test*, **Figure 3G**). Moreover, the average microglial spike amplitude per process was enhanced in microglia co-cultured with Nav1.2-L1342P neurons as well (WT+M: 0.087 ± 0.005, n=250 cells, L1342P+M: 0.121 ± 0.009, n=183 cells, \*\*\*\**p<0.0001*, *Mann-Whitney U test*, **Figure 3H**). Our human cell results suggest that diseased hyperexcitable neurons would trigger an increase in calcium signaling within the microglial processes but were insufficient to alter calcium activity in the microglial soma, surprisingly similar to the phenomena previously described in mice (Umpierre et al., 2020).

### The repetitive firing of hiPSC-derived neurons carrying the Nav1.2-L1342P is reduced with microglia co-culture

Altered microglia morphology and calcium signaling indicated that microglia could respond to intrinsically hyperexcitable neurons carrying the disease-causing Nav1.2-L1342P genetic mutations. Thus, we questioned whether human microglia would in turn affect the excitability of these neurons. To this end, we performed a whole-cell current clamp to study the repetitive firing of WT and Nav1.2-L1342P cortical neurons with or without microglia in the culture. To visualize the excitatory neuronal populations, we transduced the neurons with pAAV-CaMKIIa-eGFP, enabling fluorescent detection of excitatory neurons for patch-clamp recording. Subsequently, we seeded microglia on top of the neurons, creating a co-culture system. The co-culture was maintained for seven days, allowing for the establishment of proper interactions between microglia and neurons (**Figure 4A)**. Representative action potential firing traces show that WT neurons alone **(Figure 4B, left panel)** and in co-culture with microglia **(Figure 4B, right panel)** exhibited no significant change in their action potential (AP) firing trends. Quantitatively, the AP firing of control (WT) neurons co-cultured with microglia did not significantly differ from control (WT) neurons alone (*F*(1,28) = 0.5326, n.s *p* = 0.4716, WT: *n* = 12 neurons, two differentiations; WT+M: *n* = 18 neurons, two differentiations, *Repeated Measures Two-Way ANOVA*, **Figure 4D)**. Additionally, control (WT) neurons cultured with microglia had no difference in the maximum number of action potential firings compared to control (WT) neurons alone (WT: 8.250 ± 0.922, *n*=12 neurons, two differentiations; WT + M: 9.500 ± 0.5192, *n*=18 neurons, two differentiations; n.s *p=0.2139, Unpaired student’s t-test*, **Figure 4E**).

**Figure 4.**
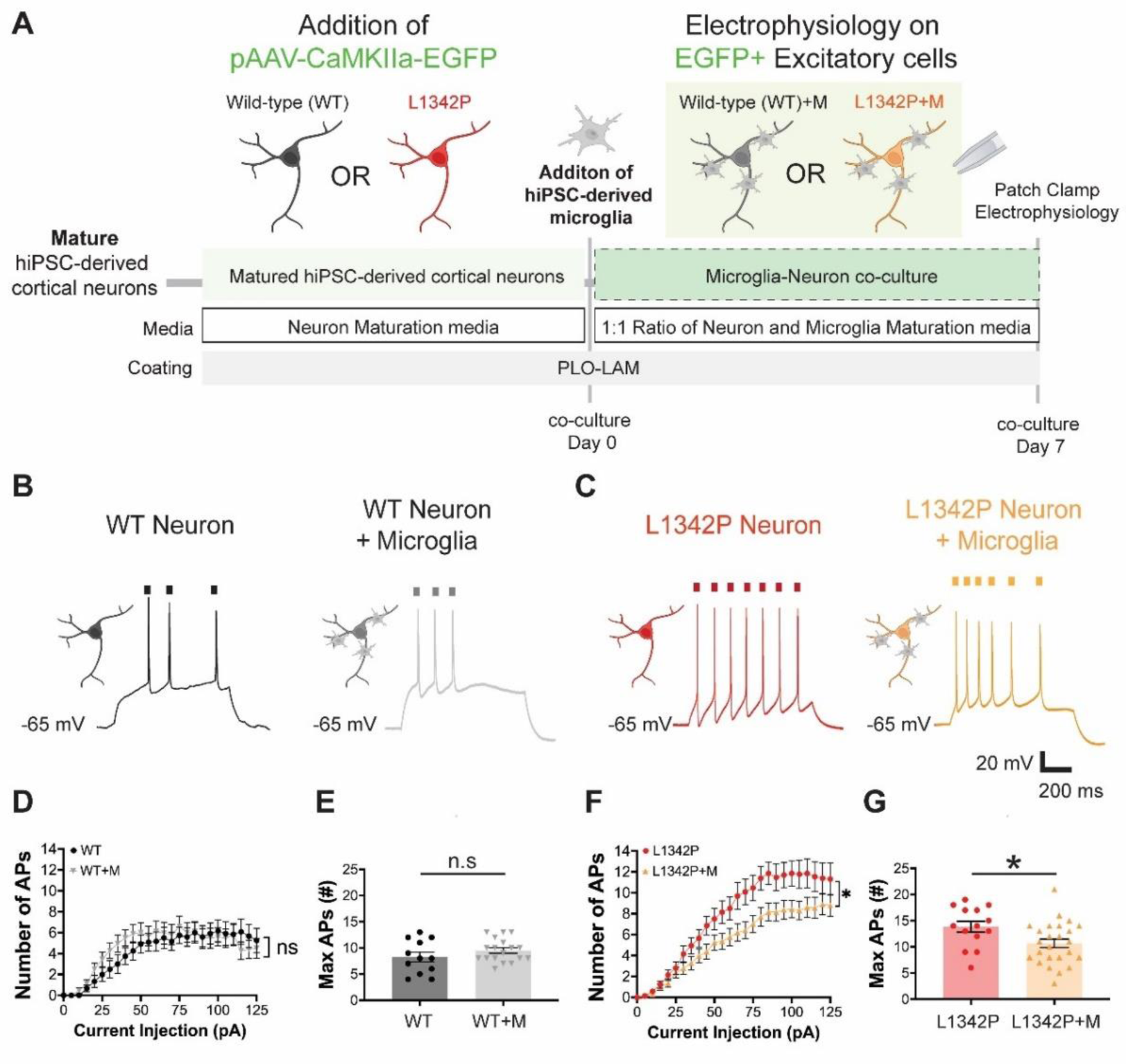
The repetitive firing of hiPSC-derived neurons carrying the Nav1.2-L1342P mutation is reduced in co-culture with microglia. **(A)** hiPSC-derived cortical neurons were transduced with an AAV-CaMKIIa-EGFP to allow the detection of excitatory neuronal populations for patch-clamp electrophysiology. Neurons and microglia were co-cultured for seven days before patch clamp measurements. **(B)** Representative action potential (AP) firings from hiPSC-derived control (WT) cortical neurons alone (left) and with microglia (right). **(C)** Representative AP firings from hiPSC-derived Nav1.2-L1342P cortical neurons alone (left) and with microglia (right). **(D)** The number of action potentials presented no statistical difference between WT neurons alone and with microglia co-culture (WT: n=12 neurons, two differentiations; WT+M: n = 18 neurons, two differentiations). (**E)** The maximum number of APs triggered from each neuron under the 0 to 125-pA current injections range presented no statistical difference between WT neurons with and without microglia co-culture (WT: n=12 neurons, two differentiations; WT+M: n = 18 neurons, two differentiations). **(F)** The action potential number of Nav1.2-L1342P neurons in co-culture with microglia consistently fired fewer APs than Nav1.2-L1342P neurons alone (L1342P: n = 14 neurons, two differentiations; L1342P+M: n = 25 neurons, two differentiations). **(G)** The maximum number of APs triggered from each neuron between the 0 to 125 pA current injections range was reduced for Nav1.2-L1342P neurons in co-culture with microglia compared to Nav1.2-L1342P neurons alone (L1342P: n*=*14 neurons, two differentiations; L1342P+M: n*=*25 neurons, two differentiations). Nav1.2-L1342P neurons co-cultured with microglia display a reduction in the maximum number of action potentials compared to Nav1.2-L1342P neurons alone. Data are presented as mean ± s.e.m. Unpaired Student’s t-test analyzed data in E and G. Each dot corresponds to one neuron. Data in D and F were analyzed by *Repeated Measures of Two-Way ANOVA*, with data pooled from at least two differentiations per condition. *p < 0.05, and n.s. (not significant).

However, a distinct finding emerged when using the Nav1.2-L1342P neurons. In isolation, the Nav1.2-L1342P neurons **(Figure 4C, left panel)** displayed extensive action potential firing at higher current injections, consistent with our previous work (Que et al., 2021). Notably, when co-cultured with microglia, the Nav1.2-L1342P neurons exhibited reduced firing **(Figure 4C, right panel)**, indicating that microglia could modulate the neuronal activity of hyperexcitable Nav1.2-L1342P neurons. Quantitatively, the Nav1.2-L1342P neurons co-cultured with microglia fired significantly fewer action potentials than Nav1.2-L1342P neurons alone (F(1,37) = 6.421, \**p* =0.0156, L1342P: n*=14* neurons, two differentiations; L1342P+M: *n* = 25 neurons, two differentiations, *Repeated Measures Two-Way ANOVA*, **Figure 4F**). Moreover, we observed a decrease in the maximum number of AP firings triggered by increasing current injections in the L1342P neuron co-cultured with microglia, showing a 21% decrease versus Nav1.2-L1342P neurons alone (L1342P: 13.860 ± 1.037, n *=*14 neurons, L1342P+M: 10.680 ± 0.812, *n*=25 neurons; \**p*=0.023, *Unpaired student’s t-test*, **Figure 4G**). Our data suggest that hiPSC-derived human microglia decreased repetitive firings in hiPSC-derived diseased neurons carrying the seizure-related Nav1.2-L1342P mutation.

### The presence of hiPSC-derived microglia reduces the maximum sodium current density in hiPSC-derived neurons carrying the Nav1.2-L1342P mutation

It is known that sodium channel density may affect the firing properties of neurons (Motipally et al., 2019). Moreover, studies have suggested that adding microglia to neurons may alter gene expression patterns (Baxter et al., 2021). Thus, the presence of microglia may lead to alternations of genes/protein levels involved in neuronal electrical activity, such as those that encode ion channels. Therefore, we performed experiments to study the sodium channel activation and maximum sodium current density in neurons cultured in the absence or presence of microglia **(Figure 5A-B)**. We obtained the sodium channel activation curve over varying voltage for Nav1.2-L1342P neurons in isolation or co-culture with microglia **(Figure 5C left).** This allowed us to measure the maximum voltage-gated sodium channel current density in Nav1.2-L1342P hiPSC-derived neurons under both conditions. Interestingly, we observed a decrease in the maximum sodium current density in Nav1.2-L1342P neurons co-cultured with microglia compared to L1342P neurons alone (L1342P: −130.8±8.6 pF/pA, n=31 neurons; L1342P+M: −101.4±6.9 pF/pA, n=28 neurons; *p=0.0107, *Unpaired Student’s t-test*, **Figure 5C, right**). This reduced maximum sodium channel current density in Nav1.2-L13242P neuron co-culture with microglia may serve as a possible mechanism underlying decreased intrinsic excitability of neurons carrying the Nav1.2-L1342P mutation when co-cultured with microglia.

**Figure 5.**
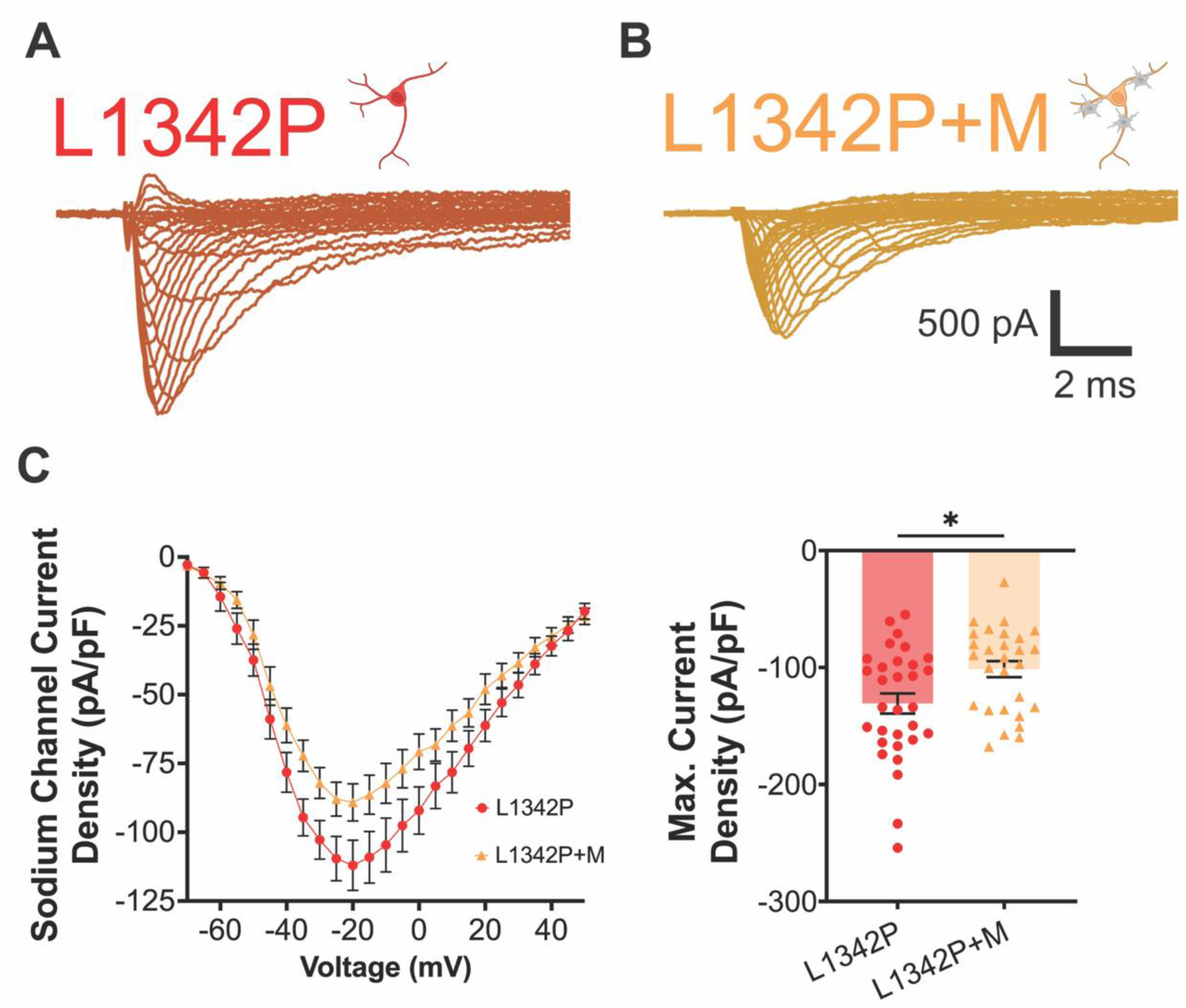
hiPSC-derived Nav1.2-L1342P neurons display reduced sodium current when co-cultured with human microglia. **(A)** Representative sodium current trace of Nav1.2-L1342P cortical neurons. **(B)** Representative sodium current trace of Nav1.2-L1342P cortical neurons in co-culture with hiPSC-derived microglia. In both A and B, the outward current was blocked using the tetraethylammonium chloride in the bath solution. **(C)** Sodium channel activation curve over varying voltage for Nav1.2-L1342P neurons in isolation or co-culture with microglia was plotted (left). The average maximum current density was significantly decreased in L1342P+M neurons compared with L1342P neuron alone (right) (L1342P: n=31 neurons, three differentiations: L1342P+M: n=28 neurons from two differentiations). Data are presented as mean ± s.e.m. Data in C right were analyzed *by Student’s t-test*. Each dot represents one neuron (right panel). Data were collected from at least two independent differentiations per genotype. *p < 0.05.

## Discussion

In this current study, we developed a co-culture model combining hiPSC-derived microglia and cortical neurons. To our knowledge, our study is the first to investigate how human microglia responds to and influence hyperexcitable neurons carrying a disease-causing mutation identified in patients with genetic epilepsy. Our findings demonstrated that microglia can sense and respond to hyperexcitable neurons by increasing branch length and calcium signaling. Interestingly, we observed that microglia significantly dampened the repetitive action potential firings of neurons carrying the epilepsy-associated Nav1.2-L1342P mutation while having minimal impact on wild-type (WT) neurons. We further showed that hiPSC-derived cortical neurons carrying the epilepsy-associated Nav1.2-L1342P mutation have reduced sodium current density when co-cultured with human microglia, which might suggest a potential mechanism by which microglia alleviate neuronal hyperexcitability. These results underscore the significance of neuron-microglia interactions in understanding disease manifestation.

Previous studies have shown that microglia can experience dynamic morphological changes in response to neuronal activity (Nebeling et al., 2023). Using a kainic acid (KA)-triggered seizure model, studies demonstrated changes in microglia morphology manifesting as an extended cell process length and increased number of microglial branch processes (Eyo et al., 2014). Surprisingly similar to previous findings, our study revealed that human iPSC-derived microglia also exhibit morphological changes in response to neuronal hyperexcitability due to genetic mutations, depicted as increased process length and microglial perimeter. These findings together suggest that microglial morphology alterations in the presence of neuronal hyperactivity are not limited to rodents and can be observed in human model system as well.

Microglial calcium signaling can undergo dynamic changes in response to their environment, including in response to the excitability of the neurons of awake mice (Umpierre et al., 2020). It was found that in response to electrical stimulation, a population of microglia endogenously expressing the Ca^2+^ indicator GCaMP6m exhibited noticeable Ca^2+^ elevations in neonatal mice (Logiacco et al., 2021). Furthermore, microglia have been observed to experience calcium fluctuations in response to changes in neuronal activity induced through pharmacological and chemogenetic methods. For example, increased neuronal activity provoked by KA-induced status epilepticus and genetic manipulations using CaMKIIa-driven Gq-DREADD activation can trigger the elevation of calcium signal amplitude and AUC in microglia (Umpierre et al., 2020). Interestingly, reduced neuronal activity through isoflurane anesthesia or CaMKIIa-driven Gi-DREADD inhibition have also been associated with increased microglial calcium activity (Umpierre et al., 2020). These observations highlight the dynamic nature of microglial responses to bimodal alterations in neuronal excitability. Notably, these calcium responses were mainly localized to microglial processes rather than the soma (Umpierre et al., 2020). To further provide insights into microglial responses to neuronal activity in a more pathophysiological-relevant context, we use an intrinsically hyperexcitable human neuronal model that does not require external stimuli to heighten neuronal activity. Although there are considerable species differences between human and rodent microglia, our study revealed a consistent finding that the microglial calcium signal was increased in the presence of hyperexcitable neurons and mainly localized in the microglial processes. While potential mechanisms and pathways (e.g., microglia sensing of excessive ATP, UDP, etc) underlying the increase in Ca^2+^ signal need to be further elucidated in follow-up studies, our current work extends the earlier observations described in rodent models to a human cell context. Together, these results indicate a potential cross-species conserved mechanism of microglial responses to neuronal excitability.

While our study found that human microglia can mitigate the excitability of hyperexcitable neurons carrying the epilepsy-associated Nav1.2-L1432P mutation, the influences of microglia on neuronal excitability and seizure phenotypes are quite complex and probably context-dependent. Microglial activation, triggered by neuroinflammation derived from trauma, stroke, febrile seizures, status epilepticus, infections, and genetic mutations, can be pivotal in epileptogenesis (Vezzani et al., 2013). In rodent models of seizures, activated microglia exerted pro-convulsive effects by releasing pro-inflammatory mediator cytokines, which can contribute to severe status epilepticus (SE) (De Simoni et al., 2000; Libbey et al., 2011; Wu et al., 2020; Henning et al., 2023). Similar findings can also be observed in cases of human temporal lobe epilepsy, further highlighting the role of inflammation in seizure formation (Kan et al., 2012). These studies together suggest a possible detrimental role of microglia as a seizure trigger. In other cases, however, microglia seem to play a beneficial role in limiting seizures and preventing pathological phenotypes. This notion is supported by experiments where the targeted removal of microglia using pharmacological (CSF1R inhibitor, PLX3397) and through genetic approaches resulted in increased neuronal activity and seizures (Szalay et al., 2016), while their repopulation conferred seizure protection (Wu et al., 2020; Gibbs-Shelton et al., 2023). In addition, in a P2RY12 Knockout (KO) mouse model subjected to KA-induced seizures, the lack of microglial process extensions worsened seizure outcomes, suggesting that microglial process extensions can be neuroprotective (Eyo et al., 2014). Moreover, the presence of microglia is believed to limit excessive neuronal activity triggered by chemoconvulsant agents in mice (Badimon et al., 2020). In our human cell-based model, we observed that the presence of microglia could reduce the excitability of the hyperexcitable Nav1.2-L1342P neurons, suggesting a presumably beneficial role of microglia in dampening neuronal hyperexcitability in this specific context. Our findings underscore the crucial role of human microglia in regulating the activity of neuronal carrying epilepsy-causing genetic mutation, highlighting the potential beneficial role of microglia in maintaining neural homeostasis. Further studies with diverse models are necessary to elucidate the precise role of microglia in different types of seizure models and to resolve the discrepancies among different studies.

While our results are revealing, they have limitations and open up additional questions. First, this is an *in vitro* study, and it is uncertain to what extent the findings could be translated into an *in vivo* context. This limitation may be addressed with human-mouse chimeric brain models we are currently establishing, in which the human iPSC-derived neurons are implanted into the brain of immunosuppressed mice for study. Moreover, as an *in vitro* study, it is also worth noting that microglia morphology in our model cannot be directly compared with microgel morphology *in* vivo (Wyatt-Johnson et al., 2017). Second, this study did not comprehensively examine the whole array of microglia functions. For example, while we identified a presumably beneficial role of microglia in our disease model, we did not assess the microglial cytokine release in our study, given that cytokine release has been shown to exacerbate neuronal hyperexcitability. Moreover, as the influence of microglia on neuronal excitability is complex, it is possible that our findings may not be generalized to other types of epilepsies (e.g., other genetic epilepsies caused by Nav1.1/Nav1.6 mutations), as microglial effects may vary depending on the content. Furthermore, our study intentionally did not utilize microglia derived from hiPSCs carrying the Nav1.2-L1342P mutation, as clear evidence suggests that Nav1.2 expression is minimal in microglia (Black et al., 2009; Black and Waxman, 2012, 2013; Abud et al., 2017; Grubman et al., 2020; Dräger et al., 2022). However, while highly unlikely, we cannot rule out that human microglia carrying the Nav1.2-L1342P might unexpectedly act differently than control (WT) microglia. Lastly, although we observed an effect of microglia on neuronal excitability and sodium current density, the molecular mechanisms underlying these findings are unclear. It remains an open question why microglia mainly affect the excitability of hyperexcitable neurons but have minimal effect on WT neurons. How microglia could modulate sodium channel density is also intriguing, with possible explanations including changes in ion channel expression, phosphorylation states, or trafficking of ion channels. While the exact mechanism is unknown, other studies have shown that co-culturing microglia with neurons in an assembloid model could influence synapse remodeling-related gene expression (Sabate-Soler et al., 2022). Future studies are needed to further dissect the mechanism underlying microglial-mediated gene/protein expression changes in neurons, furthering our understanding of microglia-neuron interactions.

In summary, our study revealed that hiPSC-derived microglia displayed altered morphology and enhanced calcium signals when co-cultured with diseased human neurons. Importantly, we found, for the first time, that microglia exerted a specific effect on the excitability of the hyperexcitable hiPSC-derived cortical neurons carrying the *SCN2A* epilepsy-related mutation Nav1.2-L1342P but had minimal impact on control (WT) neurons. Our findings underscore the significance of neuron-microglia interactions and highlight the importance of incorporating hiPSC-derived microglia in neuron-based disease models. This co-culture platform may offer a more comprehensive system for testing therapeutic interventions utilizing hiPSC-derived neurons and microglia platforms.

## Acknowledgments

This work was supported by the National Institutes of Health (NIH)-National Institute of Neurological Disorders and Stroke (NINDS) Grants: R01 NS117585 and R01 NS123154 (to Y.Y.). M.I.O.A. is supported by the Fulbright-Colciencias Scholarship Program. This work was also supported, in part, by the Indiana Clinical and Translational Sciences Institute, in part by Award Number UL1TR002529 from the National Institutes of Health, National Center for Advancing Translational Sciences, Clinical, and Translational Sciences Award. We thank Dr. Mark Estacion for the calcium imaging assay advice. We thank Dr. Shaoyou Chu from Indiana University for providing the pHrodo-myelin. We also thank the support from the Purdue University Institute for Drug Discovery (PIDD) and the Purdue Institute for Integrative Neuroscience (PIIN). The Yang Lab thanks the FamilieSCN2A foundation for the Action Potential Award and the Hodgkin-Huxley Research Award.

## Author contributions

Z.Q, M.I.O, and Y.Y. designed research; J.C.R., W.C.S. contributed unpublished reagents/analytic tools. Z.Q, M.I.O, I.C, J.Z, K.W, J.W, T.X, C.M.O, M.W, H.H, N.C, X.C, B.D, M.H and Y.Z performed research; M.I.O, Z.Q., J.C.R., R. X., A.L.B., L-J W, C.Y., and Y.Y. analyzed and interpreted data; M.I.O and Y.Y wrote the paper with input from all authors.

## References

Abud EM, Ramirez RN, Martinez ES, Healy LM, Nguyen CH, Newman SA, Yeromin AV, Scarfone VM, Marsh SE, Fimbres C (2017) iPSC-derived human microglia-like cells to study neurological diseases. Neuron 94:278–293. e279.

Badimon A, Strasburger HJ, Ayata P, Chen X, Nair A, Ikegami A, Hwang P, Chan AT, Graves SM, Uweru JO (2020) Negative feedback control of neuronal activity by microglia. Nature 586:417–423.

Baxter PS, Dando O, Emelianova K, He X, McKay S, Hardingham GE, Qiu J (2021) Microglial identity and inflammatory responses are controlled by the combined effects of neurons and astrocytes. Cell reports 34.

Black JA, Waxman SG (2012) Sodium channels and microglial function. Experimental neurology 234:302–315.

Black JA, Waxman SG (2013) Noncanonical roles of voltage-gated sodium channels. Neuron 80:280–291.

Black JA, Liu S, Waxman SG (2009) Sodium channel activity modulates multiple functions in microglia. Glia 57:1072–1081.

Chai X, Xiao Z, Zhao Q, Wang J, Ding D, Zhang J (2023) Cognitive impairment as a comorbidity of epilepsy in older adults: analysis of global and domain-specific cognition. Epileptic Disorders.

Chen R, Peng B, Zhu P, Wang Y (2023) Modulation of neuronal excitability by non-neuronal cells in physiological and pathophysiological conditions. Frontiers in Cellular Neuroscience 17:1133445.

Chen T-W, Wardill TJ, Sun Y, Pulver SR, Renninger SL, Baohan A, Schreiter ER, Kerr RA, Orger MB, Jayaraman V (2013) Ultrasensitive fluorescent proteins for imaging neuronal activity. Nature 499:295–300.

Christensen J, Dreier JW, Sun Y, Linehan C, Tomson T, Marson A, Forsgren L, Granbichler CA, Trinka E, Illiescu C (2023) Estimates of epilepsy prevalence, psychiatric co-morbidity and cost. Seizure 107:162–171.

Crawford K, Xian J, Helbig KL, Galer PD, Parthasarathy S, Lewis-Smith D, Kaufman MC, Fitch E, Ganesan S, O’Brien M (2021) Computational analysis of 10,860 phenotypic annotations in individuals with SCN2A-related disorders. Genetics in Medicine 23:1263–1272.

De Simoni MG, Perego C, Ravizza T, Moneta D, Conti M, Marchesi F, De Luigi A, Garattini S, Vezzani A (2000) Inflammatory cytokines and related genes are induced in the rat hippocampus by limbic status epilepticus. European Journal of Neuroscience 12:2623–2633.

Dräger NM, Sattler SM, Huang CT-L, Teter OM, Leng K, Hashemi SH, Hong J, Aviles G, Clelland CD, Zhan L (2022) A CRISPRi/a platform in human iPSC-derived microglia uncovers regulators of disease states. Nature neuroscience 25:1149–1162.

Epifanio R, Giorda R, Merlano MC, Zanotta N, Romaniello R, Marelli S, Russo S, Cogliati F, Bassi MT, Zucca C (2021) SCN2A Pathogenic Variants and Epilepsy: Heterogeneous Clinical, Genetic and Diagnostic Features. Brain Sciences 12:18.

Eyo UB, Peng J, Swiatkowski P, Mukherjee A, Bispo A, Wu L-J (2014) Neuronal hyperactivity recruits microglial processes via neuronal NMDA receptors and microglial P2Y12 receptors after status epilepticus. Journal of Neuroscience 34:10528–10540.

Gibbs-Shelton S, Benderoth J, Gaykema RP, Straub J, Okojie KA, Uweru JO, Lentferink DH, Rajbanshi B, Cowan MN, Patel B (2023) Microglia play beneficial roles in multiple experimental seizure models. Glia.

Grubman A, Vandekolk TH, Schröder J, Sun G, Hatwell-Humble J, Chan J, Oksanen M, Lehtonen S, Hunt C, Koistinaho JE (2020) A CX3CR1 reporter hESC line facilitates integrative analysis of in-vitro-derived microglia and improved microglia identity upon neuron-glia co-culture. Stem cell reports 14:1018–1032.

Hendrickx DA, Schuurman KG, van Draanen M, Hamann J, Huitinga I (2014) Enhanced uptake of multiple sclerosis-derived myelin by THP-1 macrophages and primary human microglia. Journal of Neuroinflammation 11:1–11.

Henning L, Antony H, Breuer A, Müller J, Seifert G, Audinat E, Singh P, Brosseron F, Heneka MT, Steinhäuser C (2023) Reactive microglia are the major source of tumor necrosis factor alpha and contribute to astrocyte dysfunction and acute seizures in experimental temporal lobe epilepsy. Glia 71:168–186.

Jairaman A, McQuade A, Granzotto A, Kang YJ, Chadarevian JP, Gandhi S, Parker I, Smith I, Cho H, Sensi SL (2022) TREM2 regulates purinergic receptor-mediated calcium signaling and motility in human iPSC-derived microglia. Elife 11:e73021.

Kan AA, de Jager W, de Wit M, Heijnen C, van Zuiden M, Ferrier C, van Rijen P, Gosselaar P, Hessel E, van Nieuwenhuizen O (2012) Protein expression profiling of inflammatory mediators in human temporal lobe epilepsy reveals co-activation of multiple chemokines and cytokines. Journal of neuroinflammation 9:1–22.

Knowles JK, Helbig I, Metcalf CS, Lubbers LS, Isom LL, Demarest S, Goldberg EM, George Jr AL, Lerche H, Weckhuysen S (2022) Precision medicine for genetic epilepsy on the horizon: recent advances, present challenges, and suggestions for continued progress. Epilepsia 63:2461–2475.

Libbey JE, Kennett NJ, Wilcox KS, White HS, Fujinami RS (2011) Interleukin-6, produced by resident cells of the central nervous system and infiltrating cells, contributes to the development of seizures following viral infection. Journal of virology 85:6913–6922.

Liu M, Jiang L, Wen M, Ke Y, Tong X, Huang W, Chen R (2020) Microglia depletion exacerbates acute seizures and hippocampal neuronal degeneration in mouse models of epilepsy. American Journal of Physiology-Cell Physiology 319:C605–C610.

Logiacco F, Xia P, Georgiev SV, Franconi C, Chang Y-J, Ugursu B, Sporbert A, Kühn R, Kettenmann H, Semtner M (2021) Microglia sense neuronal activity via GABA in the early postnatal hippocampus. Cell Reports 37:110128.

Lukens JR, Eyo UB (2022) Microglia and neurodevelopmental disorders. Annual Review of Neuroscience 45:425–445.

McQuade A, Blurton-Jones M (2021) Human induced pluripotent stem cell-derived microglia (hiPSC-microglia). In: Induced Pluripotent Stem (iPS) Cells: Methods and Protocols, pp 473-482: Springer.

McQuade A, Coburn M, Tu CH, Hasselmann J, Davtyan H, Blurton-Jones M (2018) Development and validation of a simplified method to generate human microglia from pluripotent stem cells. Molecular neurodegeneration 13:1–13.

Merlini M, Rafalski VA, Ma K, Kim K-Y, Bushong EA, Rios Coronado PE, Yan Z, Mendiola AS, Sozmen EG, Ryu JK (2021) Microglial Gi-dependent dynamics regulate brain network hyperexcitability. Nature neuroscience 24:19–23.

Motipally SI, Allen KM, Williamson DK, Marsat G (2019) Differences in sodium channel densities in the apical dendrites of pyramidal cells of the electrosensory lateral line lobe. Frontiers in Neural Circuits 13:41.

Nebeling FC, Poll S, Justus LC, Steffen J, Keppler K, Mittag M, Fuhrmann M (2023) Microglial motility is modulated by neuronal activity and correlates with dendritic spine plasticity in the hippocampus of awake mice. Elife 12:e83176.

Olson JK, Miller SD (2004) Microglia initiate central nervous system innate and adaptive immune responses through multiple TLRs. The Journal of Immunology 173:3916–3924.

Oyrer J, Maljevic S, Scheffer IE, Berkovic SF, Petrou S, Reid CA (2018) Ion channels in genetic epilepsy: from genes and mechanisms to disease-targeted therapies. Pharmacological reviews 70:142–173.

Paolicelli RC, Bolasco G, Pagani F, Maggi L, Scianni M, Panzanelli P, Giustetto M, Ferreira TA, Guiducci E, Dumas L (2011) Synaptic pruning by microglia is necessary for normal brain development. science 333:1456-1458.

Que Z, Olivero-Acosta MI, Zhang J, Eaton M, Tukker AM, Chen X, Wu J, Xie J, Xiao T, Wettschurack K (2021) Hyperexcitability and pharmacological responsiveness of cortical neurons derived from human iPSCs carrying epilepsy-associated sodium channel Nav1. 2–L1342P genetic variant. Journal of Neuroscience 41:10194-10208.

Sabate-Soler S, Nickels SL, Saraiva C, Berger E, Dubonyte U, Barmpa K, Lan YJ, Kouno T, Jarazo J, Robertson G (2022) Microglia integration into human midbrain organoids leads to increased neuronal maturation and functionality. Glia 70:1267–1288.

Schafer DP, Lehrman EK, Kautzman AG, Koyama R, Mardinly AR, Yamasaki R, Ransohoff RM, Greenberg ME, Barres BA, Stevens B (2012) Microglia sculpt postnatal neural circuits in an activity and complement-dependent manner. Neuron 74:691–705.

Schindelin J, Arganda-Carreras I, Frise E, Kaynig V, Longair M, Pietzsch T, Preibisch S, Rueden C, Saalfeld S, Schmid B (2012) Fiji: an open-source platform for biological-image analysis. Nature methods 9:676-682.

Speicher AM, Wiendl H, Meuth SG, Pawlowski M (2019) Generating microglia from human pluripotent stem cells: novel in vitro models for the study of neurodegeneration. Molecular Neurodegeneration 14:1–16.

Sun H, Li X, Guo Q, Liu S (2022) Research progress on oxidative stress regulating different types of neuronal death caused by epileptic seizures. Neurological Sciences 43:6279–6298.

Szalay G, Martinecz B, Lénárt N, Környei Z, Orsolits B, Judák L, Császár E, Fekete R, West BL, Katona G (2016) Microglia protect against brain injury and their selective elimination dysregulates neuronal network activity after stroke. Nature communications 7:11499.

Umpierre AD, Bystrom LL, Ying Y, Liu YU, Worrell G, Wu L-J (2020) Microglial calcium signaling is attuned to neuronal activity in awake mice. Elife 9:e56502.

Vezzani A, Friedman A, Dingledine RJ (2013) The role of inflammation in epileptogenesis. Neuropharmacology 69:16–24.

Weinhard L, Di Bartolomei G, Bolasco G, Machado P, Schieber NL, Neniskyte U, Exiga M, Vadisiute A, Raggioli A, Schertel A (2018) Microglia remodel synapses by presynaptic trogocytosis and spine head filopodia induction. Nature communications 9:1228.

Whitney R, Sharma S, Ramachandrannair R (2023) Sudden unexpected death in epilepsy in children. Developmental Medicine & Child Neurology.

Wolff M, Johannesen KM, Hedrich UB, Masnada S, Rubboli G, Gardella E, Lesca G, Ville D, Milh M, Villard L (2017) Genetic and phenotypic heterogeneity suggest therapeutic implications in SCN2A-related disorders. Brain 140:1316–1336.

Wu W, Li Y, Wei Y, Bosco DB, Xie M, Zhao M-G, Richardson JR, Wu L-J (2020) Microglial depletion aggravates the severity of acute and chronic seizures in mice. Brain, behavior, and immunity 89:245–255.

Wyatt-Johnson SK, Herr SA, Brewster AL (2017) Status epilepticus triggers time-dependent alterations in microglia abundance and morphological phenotypes in the hippocampus. Frontiers in Neurology 8:700.

Yang X-R, Ginjupalli VKM, Theriault O, Poulin H, Appendino JP, Au PYB, Chahine M (2022) SCN2A-related epilepsy of infancy with migrating focal seizures: Report of a variant with apparent gain- and loss-of-function effects. Journal of Neurophysiology 127:1388–1397.

Yokoi T, Enomoto Y, Tsurusaki Y, Naruto T, Kurosawa K (2018) Nonsyndromic intellectual disability with novel heterozygous SCN2A mutation and epilepsy. Human Genome Variation 5:20.

Zeng Q, Yang Y, Duan J, Niu X, Chen Y, Wang D, Zhang J, Chen J, Yang X, Li J (2022) SCN2A-Related Epilepsy: The Phenotypic Spectrum, Treatment and Prognosis. Frontiers in Molecular Neuroscience 15.

